# GNAT family Pat2 lysine acetylation of glycerol kinase and its key role in glycerol metabolism in hypersaline-adapted archaea

**DOI:** 10.1101/2025.05.27.656527

**Authors:** Heather N. Judd, Karol M. Sanchez, Leah S. Dublino, Gabriel J. Zhang, Daniel Gal, Ricardo L. Couto-Rodríguez, Julie A. Maupin-Furlow

## Abstract

Lysine acetylation is a widespread post-translational modification (PTM) involved in regulating key biological processes including central metabolism and chromatin dynamics, yet its roles in archaea remain poorly understood. Here, we investigated two GNAT (Gcn5-related N-acetyltransferase) family homologs, *pat1* and *pat2*, in the halophilic archaeon *Haloferax volcanii* (Hv). We found that a *pat2* mutant exhibited impaired growth and premature cell death on glycerol, a phenotype not observed in the parent strain, *pat1* mutant, or during growth on glucose. Complementation with plasmid-expressed *pat2* restored growth on glycerol, confirming this biological role. *In vitro* assays demonstrated that HvPat2 catalyzes the lysine acetylation of HvGlpK, a glycerol kinase essential for glycerol metabolism. Computational modeling predicted that HvPat2 residues E105, Y154, V110, and N147 may form hydrogen bonds with acetyl-CoA. To assess the functional importance of these residues, alanine substitutions were introduced at each site. Growth assays revealed that E105A and Y154A variants failed to restore growth on glycerol, while V110A and N147A had no significant effect. *In vitro*, HvPat2 Y154A, E105A, and V110A lacked acetyltransferase activity toward GlpK, whereas N147A retained partial activity. HvPat2 Y154A co-purified with a protein partner, potentially explaining the discrepancy between *in vivo* and *in vitro* results. These findings highlight the critical role of the GNAT HvPat2 in mediating lysine acetylation in regulating glycerol metabolism in archaea and offer mechanistic insight into GNAT family acetyltransferases.

**IMPORTANCE:** GNAT family homologs are widespread and diverse in their use of acyl-CoAs to acylate small molecules and proteins. These functions can be difficult to predict based on *in silico* analysis alone. Here we reveal a critical role for lysine acetylation in archaeal central carbon metabolism, identifying the GNAT family homolog Pat2 as an essential regulator of glycerol utilization in *Haloferax volcanii*. The findings expand our understanding of GNAT family acetyltransferases and highlight conserved mechanisms of metabolic control by PTM across domains of life.

## INTRODUCTION

Lysine acetylation (Kac) is an ancient form of post-translational modification (PTM) common to all domains of life that regulates metabolism, chromatin remodeling, transcription, cell structure and other important biological functions (1, 2). Fundamental to Kac is the addition of an acetyl group to the epsilon amino (*N*^ɛ^) group of lysine residues which neutralizes a positive charge. This covalent modification can impact DNA binding, protein-protein interactions, enzyme activity, substrate binding, and protein stability (3, 4). Kac occurs spontaneously through pools of reactive acetyl-CoA and acetyl-P or enzymatically through lysine acetyl transferases (KATs). Acetyl groups are removed from lysine residues by NAD^+^-dependent sirtuins (SIRTs; silent information regulators) and Zn^2+^-dependent lysine deacetylases (KDACs or HDACs)(5, 6).

KATs cluster to the GCN5-related N-acetyltransferase (GNAT) superfamily (7–9). GNATs are diverse and widespread (10) with representative members including not only KATs but also enzymes that catalyze the N-terminal (N^α^-) acetylation of proteins (11), lysine succinylation (12), the covalent modification of non-protein substrates such as aminoglycoside antibiotics (1–3) and other activities. One of the first KATs identified was the *Salmonella enterica* Pat shown to acetylate lysine residues of acetyl-CoA synthase (ACS), a central enzyme of acetate metabolism (4). Similar KAT-ACS connections are observed in *Escherichia coli*, *Bacillus subtilis*, *Rhodopseudomonas palustris*, *Streptomyces coelicolor*, *Mycobacterium smegmatis*, and *M. tuberculosis* (5–9). Many of the bacterial KAT representatives have regulatory domain(s) fused to the GNAT catalytic domain (13). The *E. coli* PatZ and *Salmonella enterica* SePat are examples of KATs fused to a nucleoside diphosphate (NDP)-forming CoA ligase-related domain (6). Likewise, *Mycobacterium tuberculosis* bKAT has an allosteric cyclic AMP (cAMP)-binding domain (14), and *Micromonospora aurantiaca* bKAT features an amino acid-sensing domain (15). In yeast, many GNATs exist with regulatory domains or as multisubunit complexes (16, 17). By comparison, archaeal GNATs are typically single-domain proteins (Pfam PF00583).

While diverse, GNATs have a common catalytic core. This core includes a classical secondary structure consisting of six to seven β-strands and four to five α-helices where a β-bulge is formed around the β4-strand and the acetyl-CoA binding site is formed by a V-like cleft between the β4 and β5 strands (18). Acetyl-CoA binding is facilitated by the phosphate binding loop positioned around β4 and α3, called the P-loop, which is formed by a conserved set of residues (Q/R-x-x-G-x-G/A) where x is any amino acid, and the nitrogenous backbone stabilizes the pyrophosphate moiety of coenzyme A (18).

Given their distinct phylogenetic relationship to bacteria and eukaryotes, archaea serve as excellent models for studying the evolution of protein function; however, biochemical studies on archaeal members of the GNAT superfamily remain limited. Although a 2Fe-2S ferredoxin of *Halobacterium salinarum* is one of the first reports of a lysine acetylated protein in prokaryotes (19), the KAT catalyzing this Kac remains to be determined. Instead, much of the current knowledge of archaeal GNATs and lysine acetylation is from the work of *Sulfolobus solfataricus* SsPat that acetylates Alba at K16, decreasing the nucleic acid binding affinity of this chromatin remodeling protein (20, 21). In *Haloferax volcanii,* HvPat1 (HVO_1756), HvPat2 (HVO_1821), and Elp3 (HVO_2888) are identified as GNAT homologs (22). These *H. volcanii* GNATs have limited primary amino acid sequence homology to the SsPat. In a proteomics study, the abundance of HvPat1 and HvPat2 are found inversely correlated with HvPat1 being downregulated during hypochlorite stress whereas HvPat2 is upregulated (23).

In this study, we investigated the function of two GNAT family lysine acetyltransferases, HvPat1 and HvPat2, in the model archaeon *Haloferax volcanii*. We focus on understanding the role of these enzymes in central carbon metabolism, particularly during growth on glycerol. Through genetic, biochemical, and structural analyses, we identify Pat2 as a key regulator of glycerol metabolism, acting via acetylation of the glycerol kinase GlpK. Our findings provide new insights into the functional significance of lysine acetylation in archaea and broaden our understanding of GNAT family enzymes in metabolic regulation.

## EXPERIMENTAL PROCEDURES

### Materials

Biochemicals were purchased from Fisher Scientific (Atlanta, GA, USA), Bio-Rad (Hercules, CA, USA), and Sigma-Aldrich (St. Louis, MO, USA). Oligonucleotides and DNA Sanger sequencing services were purchased from Eurofins Genomics (Louisville, KY, USA). Phusion High-Fidelity DNA Polymerase and other DNA modification enzymes were purchased from New England Biolabs (NEB) (Ipswich, MA, USA). DNA fragments were isolated using NEB Monarch PCR & DNA Cleanup Kit and DNA Gel Extraction Kit (Ipswich, MA, USA). Acetyl-CoA (cat. No. 16160) was purchased from Cayman Chemical (Ann Arbor, Michigan, USA).

### Strains, media, and growth conditions

Strains used in this study are summarized in **Table S1**. *E. coli* cultures were grown at 37°C in Luria-Bertani (LB) medium supplemented with ampicillin (100 µg/mL) (Amp) and/or chloramphenicol (34 µg/mL) (Cam) as needed. *H. volcanii* strains were grown at 42 °C in ATCC974, Hv-Ca+ or minimal media as previously described with the following modifications (24). Minimal medium was composed of 18% salt-water, 30 mM Tris-Cl, pH 7.5,3 mM CaCl_2_, 10 mM NH_4_Cl, 3 mM of desired carbon source (glycerol, fructose, or glucose), 0.975 mM KPO_4_ and 50 µg/mL uracil along with trace elements, thiamine, and biotin at concentrations as previously described (24). *H. volcanii* medium was supplemented with novobiocin (0.2 µg/mL), uracil (50 µg/mL), and/or tryptophan (2 mM) as needed. Liquid cultures were grown by angled rotation in culture tubes (13 × 100 mm^2^) (Glas-Col mini-rotator, Terre Haute, IN) or continuous orbital shaking at 200 rpm (New Brunswick Scientific I 24 Incubator Shaker, Edison, NJ) as indicated. Culture plates were supplemented with 1.5% (w/v) agar. Growth was measured by optical density at 600 nm (OD_600_) where 1 OD_600_ unit equals approximately 10^9^ CFU/mL. All strains, including those carrying plasmids, were stored in 20% (v/v) glycerol stocks at -80°C until use.

### Strain and plasmid constructions

Plasmids and primers synthesized and used in this study are summarized in **Table S1**. For all plasmid constructions, DNA fragments for cloning were generated by PCR using high-fidelity Phusion DNA polymerase. The genomic DNA used for PCR template was extracted from *H. volcanii* H26 following the protocol outlined in *The Halohandbook* (24). PCR products were purified using the Monarch PCR and DNA Cleanup Kit (New England Biolabs). PCR products were digested with restriction enzymes (as indicated in **Table S1**) according to the manufacturer’s protocols. After digestion, the DNA fragments were cleaned using PCR clean-up (New England Biolabs) and ligated using Quick T4 DNA ligase at room temperature (RT) for 15 min. DNA ligation products were transformed into *E. coli* Top10. Plasmid DNA was extracted into DNase/RNase-free deionized water (ThermoFisher) using the PureLink Miniprep Kit (Invitrogen). PCR was used for screening and Sanger DNA sequencing was used to verify the fidelity of the plasmid insert (Eurofins Genomics, Louisville, KY).

Plasmids were transformed into the *dam^−^ dcm^−^* strain *E. coli* GM2163 to remove methylation of the DNA prior to transformation of *H. volcanii* strains according to the *Halohandbook* (24). Selection markers for *H. volcanii* included *gyrB* (encoding a modified GyrB subunit of DNA gyrase rendering cells resistant to novobiocin) and *pyrE2* (encoding orotate phosphoribosyl transferase rendering *ΔpyrE2* strains viable in Hv-Ca+ medium devoid of uracil).

To generate the *pat2-strepII* expression plasmids (encoding HvPat2 fused to a C-terminal StrepII tag), the *pat2* gene was PCR amplified from *H. volcanii* genomic DNA using primers HVO_1821_NdeI and HVO_1821-KpnI. The PCR product was ligated into the NdeI and KpnI sites of pJAM809 to generate plasmid pJAM4017 carrying the *pat2-strepII* gene under control of the P2*_rrnA_* promoter. The *pat2-strepII* gene was amplified from pJAM4017 by PCR using Pat2_ecori_F and Pat1_rev primers. The PCR product was ligated into the NdeI and BamHI sites of pTA963 to generate plasmid pJAM4554, carrying the *pat2-strepII* gene under control of the tryptophan-inducible promoter P*_tnaA_*.

Site-directed mutations (SDMs) of *pat2-strepII* were generated on pJAM4017 using pat2_SDM-anchor1, pat2_SDM-anchor2, E105A_rev, V110A_rev, N147A_rev, and Y154A_rev primers using the rrPCR method as previously described(25). The resulting plasmids included pJAM4555 (Pat2-StrepII E105A), pJAM4556 (Pat2-StrepII V110A), pJAM4557 (Pat2-StrepII N147A), and pJAM4558 (Pat2-StrepII Y154A). All *pat2-strepII* wild-type and SDM plasmids were transformed into *H. volcanii* strain JM502 (H26 *Δpat2*).

The *glpK* gene was isolated from *H. volcanii* genomic DNA by PCR using primers F-glpK and R-glpK. The 1.5 kb PCR fragment was digested, purified, and ligated into the NdeI and BlpI sites of plasmid pET15b. The resulting plasmid, pJAM4360 (pET15b expressing *Hv*GlpK), was transformed in *E. coli* Top10 for storage in the 20 % (v/v) glycerol stocks at -80 °C.

### Growth Curves

Strains were streaked from the 20 % (v/v) glycerol stocks onto ATCC974 plates and incubated for ∼7 days at 42°C. Novobiocin was included in the medium for strains carrying the pJAM202c empty vector and *pat2-strepII* expression plasmids (pJAM4017, pJAM4554, pJAM4556, pJAM4557, and pJAM4558). A single isolated colony was used to inoculate 5 mL of minimal medium supplemented with the designated carbon source and cultured for 2 days in rotating culture tubes at 42°C. Cells were then diluted to an optical density (OD_600_) of 0.02 in 5 mL of fresh medium and incubated again at 42°C overnight in rotating culture tubes. The following morning, cells were diluted to an OD_600_ of 0.02 again in 5 mL fresh medium and then rotated in the culture tubes at 42°C for 15 min for mixing. Cells were added to a sterile polystyrene tissue cultured treated 96-well, flat bottom cell culture plate (reproducible results with either CellPro, Alkali Scientific, Fort Lauderdale, FL or GenClone, Genesee Scientific, El Cajon, CA) where each well had a final volume of 200 µL (5 replicates per strain per media) with outer wells containing 300 µL of sterile diH_2_O. Wells with each medium type were also included as a blank control to account for background signal. The lid was secured to plate bottom using 3M Micropore tape. Growth was monitored by measuring OD_600_ every 15 min for 120 h at 42°C with aeration (continuous orbital shaking) using a microtiter plate reader (BioTek Epoch 2, Agilent Technologies, Santa Clara, CA).

### Large scale cultivation of *H. volcanii* strains for Pat2 purification

Pat2 and variant proteins were purified from *H. volcanii* H26 Δ*pat2* strains carrying the *pat2-strepII* and SDM expression plasmids. For strains carrying plasmids with the *pyrE2*^+^ marker (*e.g.,* pJAM4554), cells were freshly streaked from glycerol stocks on to Hv-Ca+ agar plates. A single colony was used to inoculate 100 mL Hv-Ca+ medium and incubated for ∼2 days at 42°C with orbital shaking in 500 mL Erlenmeyer flasks. The starter culture was subcultured (10 mL) into 500 mL of Hv-Ca+ and incubated at 42°C with orbital shaking in 2.8 L Fernbach flasks until OD_600_ reached 0.6 where expression of *pat2-strepII* ORF was induced by addition of 2 mM L-tryptophan and cultured for an additional 12-18 h. Once the desired OD_600_ of 1.2-1.4 was reached, cells were harvested by centrifugation (2,995 g x at 20°C for 50 min). Cell pellets were stored at -80°C until use. For strains carrying plasmids with the *gyrB* (Nov^R^) marker (*e.g.,* pJAM4017 and derivatives), the cells were cultured in a similar manner except ATCC974 supplemented with 0.3 µg/mL novobiocin was used for selection instead of Hv-Ca+ and no L-tryptophan was needed for induction of the *pat2-strepII* and SDMs.

### Pat2 purification

Cell pellet (usually 3 g wet weight) was resuspended in 15 mL lysis buffer (50 mM HEPES, pH 7.5, 2 M NaCl, 1 mM TCEP, 3 mM MgCl_2_, 1 mM CaCl_2_, 10 µg/mL DNase I, and Mini cOmplete EDTA-free protease inhibitor tablets (used according to manufacturer’s recommendations). Cells were disrupted using a French pressure cell with three to four passes (1,500 psi, minimum high ratio of 140, Glen-Mills, NJ, U.S.). Lysate was clarified by centrifugation (13,177 *x g* at 4°C for 30 min) and filtration (0.45 µm and 0.22 µm, PES filters, CellPro). Samples were incubated with 0.5 mL (1 mL 50 % slurry) Strep-Tactin Superflow Plus resin (Qiagen, Germantown, MD) in 10 mL columns (Pierce, ThermoFisher Scientific, U.S., Cat. No. 89898) for at least 1 h at 4°C with gentle rocking. Prior to sample application resin was equilibrated in wash buffer (50 mM HEPES, pH 7.5, 2 M NaCl, 1 mM TCEP). After application of sample, non-specific proteins were removed by subsequent rounds of washing with the wash buffer (5 washes with 2.5 mL wash buffer). Pat2-StrepII and variant proteins were eluted from the column by incubating with 300 µL wash buffer supplemented with 1 mM desthiobiotin for 30 min at 4°C, two times total for a final volume of 600 µL. Final elutions (600 µL total) were dialyzed at 4°C overnight in dialysis buffer (50 mM HEPES, pH 7.5, 2 M NaCl, 1 mM DL-1,4-dithiothreitol (Thermo Fisher, USA) using D-Tube dialyzer midi, MWCO 3.5 kDa (EMD Millipore Corporation, Burlington, MA, USA). D-tubes were prepared according to manufacturer’s directions prior to use. Pat2 fractions were combined and concentrated to ≤ 500 µL using Sartorius Vivaspin Turbo 5 10,000 MWCO centrifugal concentrator columns according to manufacturer’s directions (Fisher Scientific, USA) and then further purified by size-exclusion chromatography (SEC) using a Superdex 75 Increase 10/300 GL FPLC column (Cytiva, Marlborough, MA). Aliquots of protein samples (0.5 mL) were applied to the column equilibrated in freshly prepared filtered dialysis buffer (see above) at a flow rate of 0.3 mL/min. Fractions (0.5 mL) were collected and analyzed for Pat2 protein. The final fractions were combined and concentrated using a Vivaspin centrifugal concentrator (3 MWCO, MilliporeSigma Burlington, MA).

### Protein quantification and SDS-PAGE

Protein concentration was measured by Bradford assay with bovine serum albumin as the standard according to manufacturer’s protocol (Bio-Rad, Hercules, CA). For each measurement, sample (5 μL) was mixed with 250 μL of Bradford reagent, and the mixture was incubated for 5 min at RT. Absorbance at 595 nm (A595) was recorded using a BioTek Epoch 2 microtiter plate reader. The assay exhibited linearity within the 0 to 2,000 μg·mL⁻¹ protein range. For SDS-PAGE analysis, proteins were mixed with 2X SDS-PAGE loading buffer: 100 mM Tris-HCl buffer, pH 6.8, 4% (w/v) SDS, 20% (v/v) glycerol, 0.6 mg/mL bromophenol blue, and 5% (v/v) β-mercaptoethanol. Samples were boiled for 5-10 min and separated by reducing 12% SDS-PAGE in Tris-glycine-SDS (TGS) buffer. Precision Plus Protein Kaleidoscope molecular mass marker (BioRad, Hercules, CA) was used as the standard. Proteins were stained in gel with Coomassie Brilliant Blue R-250 for 1 h at RT and then destained with diH_2_O overnight. Gels were imaged using an iBright Imaging System (Invitrogen, Carlsbad, CA) according to the manufacturer’s protocol.

### Immunoblotting analysis

Proteins were transferred from SDS-PAGE gels to PVDF membranes (0.45 µm) (Immobilon-FL, Millipore, Burlington, MA) using the wet transfer method at 30 V for 14 h at 4°C with constant stirring as per standard protocol (BioRad, Hercules, CA, USA). After protein transfer, the membranes were incubated with gentle rocking for 2 h at RT in blocking buffer composed of TBST (0.05 M Tris-HCl, pH 7.6, 0.15 M NaCl, 0.1% (v/v) Tween 20) supplemented with 5% (w/v) skim milk powder for immunoblotting analysis of His-tagged and StrepII-tagged proteins. To detect His-tagged proteins, the membranes were incubated at RT for 1 h with a 1:10,000 dilution of HRP-conjugated mouse monoclonal anti-6×His-tag antibody (Cat. No. HRP-660005). After incubation, the membranes were washed 5 x 5 min with TBST. To detect StrepII-tagged proteins, membranes were incubated at RT for 1 h with polyclonal anti-StrepII tag antibody (NWSHPQFEK Genescript A002026 rabbit) at 0.25 µg per mL blocking buffer. After washing the membrane 5 x 5 min with TBST, the membrane was incubated for 1 h at RT with 1:10,000 dilution of goat anti-rabbit HRP-conjugated (Southern Biotech #4010-05). The membrane was similarly washed 5 x 5 min with TBST. To detect lysine acetylation, membranes were blocked overnight at 4°C in TBST supplemented with 5% (w/v) BSA and probed with anti-acetyllysine rabbit mAb (PTM Bio, #PTM-105RM) as the primary at 1:5,000 dilution and mouse anti-rabbit IgG-HRP (Santa Cruz Biotechnology; #sc-2357) as the secondary antibody at 1:10,000 dilution. Chemiluminescent signals were detected using Chemiluminescent solution according to supplier (Thermo Scientific, #89880) on an iBright FL1000 Imaging System (Thermo Fisher Scientific).

### HvGlpK purification from recombinant *E. coli*

*E. coli* Rosetta (DE3) was freshly transformed with the His-HvGlpK expression plasmid pJAM4360 and selected on LB Amp/Cam plates. Isolated colonies were used to inoculate a starter culture in 20 mL LB Amp/Cam (250 mL Erlenmeyer flask). The starter culture was grown at 37°C until an OD_600_ of 0.8 was reached. Subsequently, 500 mL large-scale cultures were inoculated to an initial OD_600_ of 0.01 and incubated at 37°C with orbital shaking. When the cultures reached an OD_600_ of 0.6–0.8, the temperature was reduced to 25°C. Once the cultures reached 25°C, protein expression was induced by adding isopropyl-β-D-1-thiogalactopyranoside (IPTG) to a final concentration of 0.4 mM, followed by overnight incubation with orbital shaking. After 16 h, cells were harvested, and pellets were either stored at -80°C or immediately resuspended in lysis buffer (50 mM HEPES, pH 7.5, 50 mM NaCl, 5 mM β-mercaptoethanol, 40 mM imidazole, 10 µg/ mL DNase, and protease inhibitors) at a ratio of 5 mL per gram of cells. IPTG was added only after the cells reached 25°C to maintain protein solubility, and pellets were resuspended and lysed in low-salt buffer before buffer exchange to high-salt conditions (2 M NaCl) using Zeba spin desalting columns 7K MWCO according to the manufacturer’s buffer exchange protocol (Thermo Scientific, Rockford, IL). This step was introduced to minimize aggregation of the ‘salt-loving’ GlpK. Ni(II)-NTA His-Bind resin (0.25 mL of 50% slurry; EMD Millipore Corporation, Burlington, Ma, USA) was equilibrated in wash buffer composed of 50 mM HEPES, pH 7.5, 2 M NaCl, 5 mM β-mercaptoethanol, and 40 mM imidazole. Cell lysate (0.5 mL) was incubated with the pre-equilibrated resin for 1 h at 4°C with gentle rocking. The resin with sample was washed three times with wash buffer (50 mM HEPES, pH 7.5, 2 M NaCl, 40 mM imidazole) and eluted two times with 200 µL elution buffer (50 mM HEPES, pH 7.5, 2 M NaCl, 250 mM imidazole) for a final elution volume of 400 µL. Dialysis was performed as previously described for Strep-II tag purifications, except 10% (w/v) glycerol is included in the HvGlpK dialysis buffer.

### Lysine acetylation activity assay

Lysine acetylation reactions (50 µL) contained: 6 µM enzyme (HvPat2), 2 µM substrate protein (HvGlpK_Ec_), 0.2 mM acetyl-CoA, 50 mM HEPES, pH 7.5, 2 M NaCl, and 1 mM DTT. Reactions were incubated at 37°C for 3 h followed by precipitation overnight on ice with 10% (w/v) trichloroacetic acid (TCA). Samples were centrifuged (10 min, 17,999 × *g*, 4°C), and protein pellets were washed with 100% cold acetone at 4°C. Air-dried protein pellets were resuspended in 20 µL 2X SDS-PAGE reducing buffer, boiled for 5 min, and resolved by 12% SDS-PAGE for Coomassie blue staining and immunoblotting as previously described.

### *In silico* modeling of HvPat2 and docking of acetyl-CoA and HvGlpK

Residues involved in binding acetyl-CoA were identified by first finding GNATs with structural homology to HvPat2 (PDBs/Accession numbers: 4NXY, AAG08251.1, AEW6158.1, AN032847.1, 2I79). These GNATs were identified using AlphaFold Foldseek (https://search.foldseek.com/search). Clustal Omega (26) was used to align the primary sequences. The resulting .aln file and the predicted Pat2 structure were imported to ESPript 3.0 webserver (https://espript.ibcp.fr/ESPript/ESPript/) (27) to align the crystal and predicted structures using HvPat2 as a reference to predict conserved secondary structure formation among the dataset. The predicted tertiary structure of HvPat2 was superimposed onto the crystal PDB structure of *S. lividans* PatA (PDB: 4NXY) bound to acetyl-CoA using the ChimeraX matchmaker structural tool (https://www.cgl.ucsf.edu/chimerax/). This overlay was used to identify residues for acetyl-CoA binding, which was later used in the docking analysis. To model molecular docking, the PDB file of the HvPat2 model was modified using ChimeraX Dock Prep. This step is necessary to ensure the enzyme is in the proper format for optimized molecular docking simulations by removing water molecules and non-protein ligands, adding necessary hydrogens, and assigning partial charges. Once prepared, the HvPat2 PDB file was uploaded to PyRx (https://pyrx.sourceforge.io/) alongside the crystal structure of acetyl-CoA (Pubchem CID: 444493). Docking was performed on the involved residues followed by energy minimization of the overall docked structure to produce the most energetically efficient binding conformation. Upon completion, the program returned models of docked acetyl-CoA: HvPat2 poses ranked based on binding energy (kcal/mol). The most energetically efficient model was chosen and visualized in ChimeraX. AlphaFold 3 was used to predict HvPat2:HvGlpK binding.

### Statistical Analysis

Growth curve and enzymatic activity assays were performed under defined conditions (as outlined in earlier sections). Each growth curve experiment included biological replicates performed in at least triplicate and was conducted a minimum of three times to ensure reproducibility. Area under the curve (AUC) values were calculated based on the trapezoidal rule and are presented as mean values ± standard deviation (SD). Statistical significance between experimental groups was assessed using Student’s t-test, with a threshold for significance set at p-value < 0.05. All statistical analyses were performed using Microsoft Excel. Enzymatic activity assay experiments were performed at least in duplicate and demonstrated to be reproducible with representative immunoblots presented.

## RESULTS

### HvPat1 and HvPat2 are GNAT homologs with conserved active site residues

*H. volcanii* (Hv) Pat1 and Pat2 are single-domain GNAT family proteins that share 32.4% amino acid sequence identity (**Figure 1**). A major distinction between these two GNATs is that HvPat1 and its close homologs have an N-terminal Cys-rich motif (C-x_5_-C-x_3_-C, where x is any amino acid) that could play a role in metal ion coordination or redox processes, but this has yet to be determined. While bacterial GNATs are often fused to regulatory domains, such as cNMP and NDP-forming acyl-CoA synthetase-like domains (13, 28–30), the single GNAT domain architecture of HvPat1 and HvPat2 is common among the archaeal homologs. GNATs, though diverse, share a conserved core including a P-loop, and many feature an active site glutamate important for catalysis (13, 28). The P-loop motif that plays a crucial role in binding the pyrophosphate group of acetyl-CoA in GNAT family homologs is conserved in HvPat1 and HvPat2, located between the predicted β5 and α3 regions based on AlphaFold 3D-structural modeling. Moreover, both *H. volcanii* homologs are predicted to be catalytically active, as they have a conserved glutamate (HvPat2 E105 and HvPat1 E119) that could serve as a general base to deprotonate the amino group of the substrate lysine residue to then facilitate nucleophilic attack on the acetyl group of acetyl-CoA during the catalytic reaction (31).

**Figure 1.**
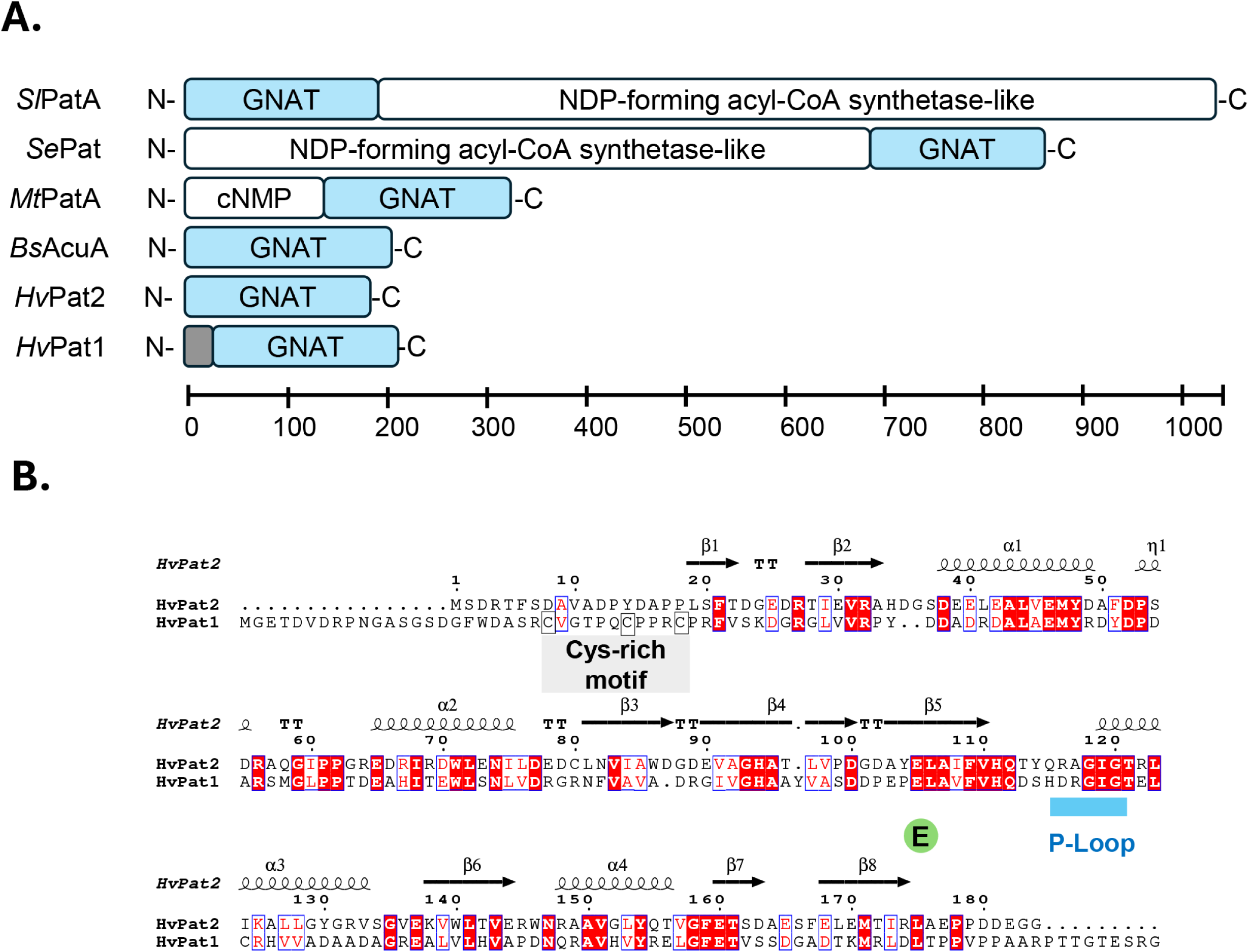
HvPat1 and HvPat2 are GNAT family homologs with conserved active site glutamate and P-loop motif. A) Domain architecture of representative GNAT family proteins. *Sl*, *Streptomyces lividans*; *Se*, *Salmonella enterica*; *Mt*, *Mycobacterium tuberculosis*; *Bs*, *Bacillus subtilis*. Grey box, conserved C-X_5_-C-X_3_-C motif of HvPat1. Modified from (30) to include archaeal proteins. B) Amino acid sequence alignment of HvPat1 and HvPat2. Conserved residues, white text and highlighted in red. Functionally similar residues, red text and boxed in blue. Highlighted: P-loop (blue), conserved CoA binding pocket located between the predicted β5 and α3 regions of HvPat2 (and HvPat1) based on AlphaFold 3D-structural modeling. P-loop plays a crucial role in binding the pyrophosphate group of acetyl-CoA in GNAT family homologs. Glu, (E green), conserved active site glutamate that may act as a general base to deprotonate the amino group of the lysine residue of the substrate during catalysis. Cys-rich motif (grey), C-x_5_-C-x_3_-C, conserved among HvPat1 homologs. Sequence alignment performed with Clustal Omega (26). Output processed by ESPript 3.0 (http://espript.ibcp.fr/ESPript/cgi-bin/ESPript.cgi)(27) .

### *Δpat2* mutants display carbon-source dependent growth defects

Post-translational modifications (PTMs), such as lysine acetylation, are crucial for cells to rapidly and dynamically adapt to shifts in carbon source utilization (32, 33). We reasoned that if HvPat1 or HvPat2 is responsible for the lysine acetylation of a key enzyme involved in carbon metabolism, then single and/or double mutations of *pat1* and *pat2* would affect growth in a carbon source-dependent manner. Therefore, *H. volcanii* parent (H26), *Δpat1*, *Δpat2*, and *Δpat1Δpat2* mutant strains were analyzed for growth in different carbon sources, including glycerol and glucose, by measuring optical density (OD_600_) over time (**Figure 2A**). All four strains were found to display similar growth curves when grown on glucose minimal media, entering exponential and stationary phases at similar times. By contrast, carbon source dependent differences were observed when the *pat2* mutant strains were examined for growth on glycerol minimal media (GMM). Thus, carbon source dependent phenotypes were observed for the *pat2* deficient strains on glycerol.

**Figure 2.**
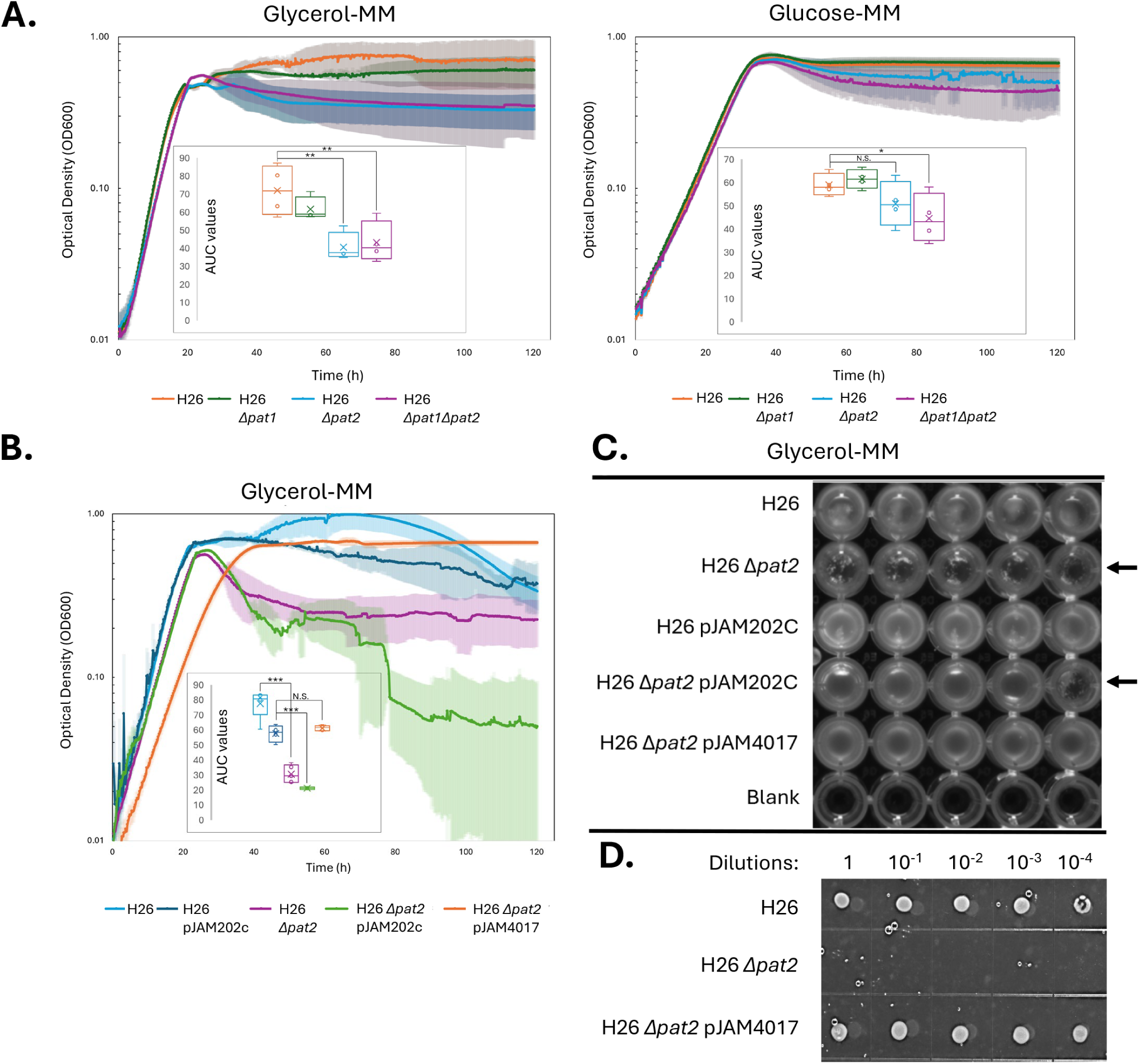
*Δpat1* and *Δpat2* mutant strains display carbon-source dependent phenotypes. **A)** Growth curves reveal carbon source-dependent growth defect of the *Δpat2* mutant on glycerol compared to glucose. H26 parent (orange), Δ*pat1* (green), Δ*pat2* (blue), and Δ*pat1* Δ*pat2* double mutant (purple). Glycerol and glucose minimal media (MM) indicated. **B)** Growth of curves demonstrate the *Δpat2* mutant can be complemented for the growth defect on glycerol by *in trans* expression of *pat2-strepII* from a plasmid. H26 (light blue), H26-pJAM202C empty vector (dark blue), H26 Δ*pat2* (purple), H26 Δ*pat2*-pJAM202C empty vector (green), and H26 Δ*pat2*-pJAM4017 *pat2*-*strepII* expression plasmid (orange) in glycerol-minimal media (GMM). The growth curves (A-B) were performed by measuring optical density (OD_600_) every 15 min for 120 h. Optical densities of each strain in each culture medium were normalized at each subculturing stage and prior to loading cells onto a 96-well plate. Control wells containing each media type (no cells) were used as a blank to subtract the background OD_600_ from each sample at each time point is presented. Each experiment was performed using at least 3 replicates and then the average OD_600_ for each strain/condition at each timepoint was taken. The blanked value for each medium type was subtracted from the averaged values and graphed in log scale. Inset graphs show area under the curve (AUC) values for each strain. * p-value ≤0.05, ** p-value <0.01, *** p-value <0.0001, N.S. no significant difference. Experiments were performed at least in duplicate and found reproducible. **C)** *Δpat2* mutants aggregate after growth in glycerol minimal medium. Imaging of plate immediately following completion of growth curve shows cell physiology. Aggregated cells indicated by arrow. Images taken using Nucleic acid setting on iBright. **D)** *Δpat2* mutants are non-viable after growth in glycerol minimal medium. Cells from the GMM liquid growth curve were spotted from serial dilutions onto GMM plates and incubated for one week at 42°C as indicated. Compared to the H26 parent and Δ*pat2* expressing *pat2-strepII* from plasmid pJAM4017, the *Δpat2* mutant was found non-viable.

To further test these findings, growth curve analyses were performed with strains that could complement the *Δpat2* mutant. For this complementation, the *Δpat2* mutant was transformed with a plasmid, pJAM4017, designed to synthesize HvPat2 with a C-terminal StrepII tag (HvPat2-StrepII). Growth defects that may be caused by harboring the plasmid were also accounted for by comparing growth of control strains carrying the empty vector plasmid, pJAM202c. These strains were examined for growth on glycerol (**Figure 2B**). This analysis reconfirmed our finding that *pat2* is important for growth on glycerol. The *Δpat2* mutant was found to be complemented for growth on glycerol when carrying the *pat2* (*pat2-strepII*) gene *in trans*. Although in early phases of curve the *Δpat2* mutant demonstrated slower growth on glycerol when carrying the *pat2* gene *in trans*, this strain achieved an overall cell yield that was comparable to the parent strain (with or without the plasmid vector). In contrast, the *Δpat2* mutant displayed a reduced cell yield on glycerol that was more pronounced when burdened with the empty vector. Further analysis by imaging (**Figure 2C**) and plating assay (**Figure 2D**) revealed that strains lacking *pat2* clumped, or aggregated in bottom of the well, and were no longer viable when compared to the parent and the *Δpat2* mutant complemented *in trans* with the *pat2*. These results suggest that expressing *pat2* (*pat2-strepII*) from the plasmid can complement the *Δpat2* mutation and that restoring *pat2* expression alleviates the drop in OD_600_ caused by the cells clumping during late stationary phase when cultured in glycerol minimal medium (GMM). These results suggest that HvPat2 likely functions as a lysine acetyltransferase and that it plays a critical role in regulating the activity of enzymes involved in glycerol metabolism.

### HvPat2 purifies as a monomer

HvPat2 fused to a C-terminal StrepII tag (HvPat2-StrepII) was purified by affinity chromatography. The protein was found to purify as single protein band that migrated at 25 kDa by SDS-PAGE (**Figure 3A-B**). Further analysis using size exclusion chromatography (SEC) revealed elution profiles consistent with HvPat2 adopting a native molecular mass that corresponds to the monomeric form **(Figure 3B inlay)**. For reference, HvPat2-StrepII tagged proteins are 196 amino acids in length with the theoretical isoelectric point (pI) calculated to be 4.2 where the estimated molecular mass is 21,636 Da. No additional protein partners were observed to co-purify with wild-type HvPat2-StrepII.

**Figure 3.**
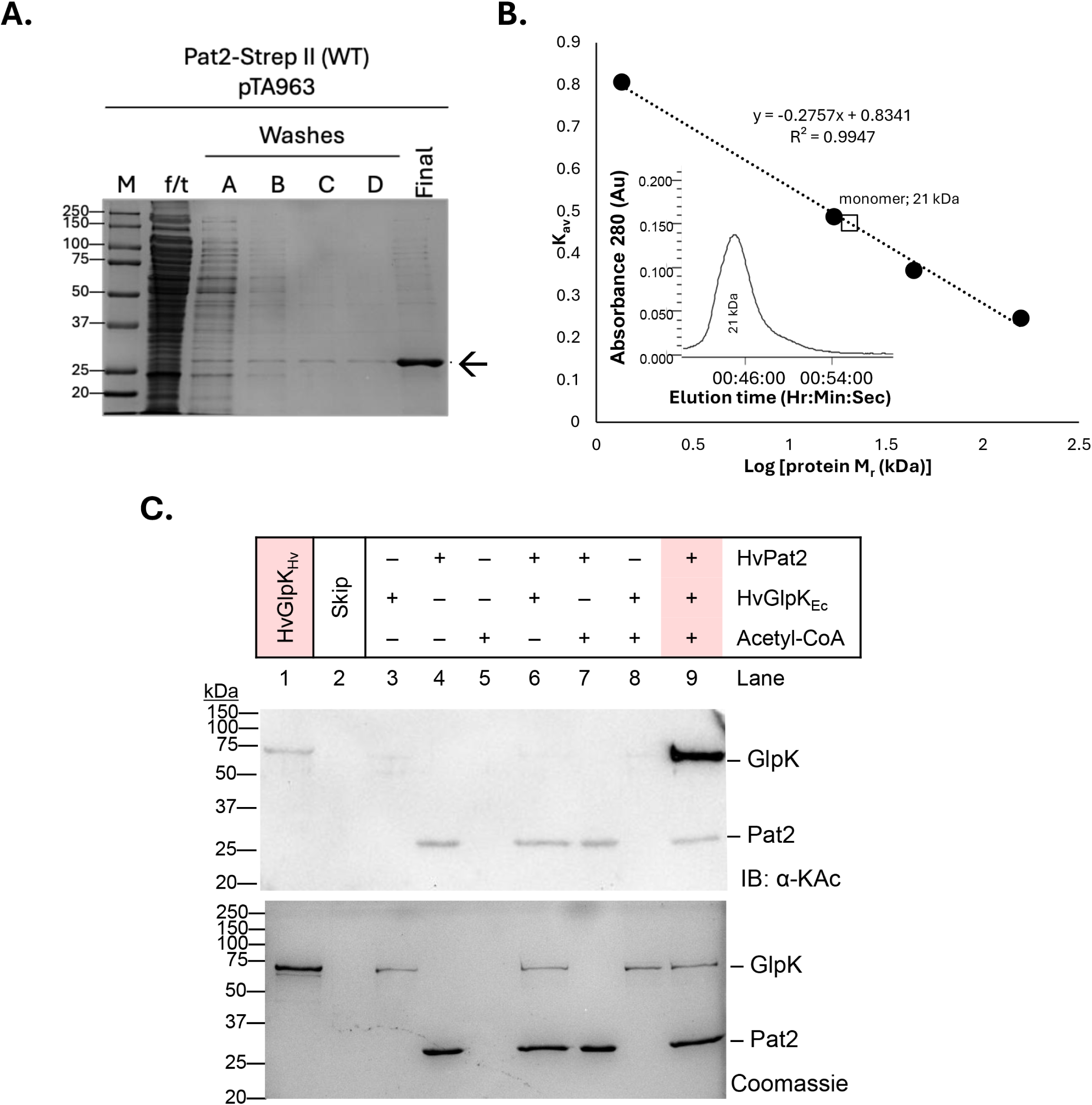
Purified HvPat2 catalyzes the lysine acetylation of glycerol kinase (HvGlpK). A) HvPat2-StrepII enriched by StrepTactin affinity chromatography. HvPat2-StrepII was purified from 3-3.5 g (wet weight) cell pellet of H26 Δ*pat2*-pJAM4554 expressing the StrepII tagged protein *in trans*. Cells were cultured to stationary phase in Hv-Ca^+^ medium. Cells were lysed with French press. Cell lysate was clarified and applied to StrepTactin resin. Flowthrough (f/t) was collected, resin was washed (A-B), and f/t was reapplied to column for a total of four washes (A-D). HvPat2-StrepII was eluted with binding buffer supplemented with 1 mM desthiobiotin. Fractions of f/t (1 µL), first set of washes (5 µL), and final eluted protein (5 µL) were separated on 12% SDS-PAGE. Protein gels were stained with Coomassie blue for 30 min, briefly rinsed with diH_2_O, then destained with fresh diH_2_O. (see materials and methods for details). B) Size exclusion chromatography demonstrates HvPat2-StrepII purifies as a monomer with a molecular mass of 21 kDa. C) HvPat2 catalyzes the lysine acetylation of HvGlpK. Reactions included HvPat2, acetyl-CoA and HvGlpK purified from recombinant *E. coli* (_EC_) as indicated. Coomassie Blue stain was used to control for protein loading. Endogenously acetylated glycerol kinase purified from *H. volcanii* (HvGlpK_Hv_) served as a positive control. Lysine acetylation reactions were assembled *in vitro* using purified HvPat2-StrepII (6 µM) as enzyme, His-HvGlpK_Ec_ (2 µM) as protein substrate, and acetyl-CoA (0.2 mM) as the acetyl group donor (materials and methods). Reactions were incubated at 37°C for 3 h. Products were resolved by 12 % SDS-PAGE and analyzed by immunoblotting using anti-acetyl-lysine (IB:αKac) and anti-His tag (IB:αHis) antibodies.

### HvPat2 catalyzes the lysine acetylation of HvGlpK

We reasoned that if HvPat2 is essential for glycerol metabolism in *H. volcanii* then it would act on key enzymes involved in processing glycerol, therefore we investigated its ability to acetylate HvGlpK, a glycerol kinase essential for the phosphorylation of glycerol to glycerol-3-phosphate (10). HvGlpK is acetylated at K153 based on our previous analysis of the lysine acetylome of cells grown on glycerol (34). To isolate non-acetylated HvGlpK, the *H. volcanii glpK* (*hvo_1541*) gene was cloned into a pET vector that expresses proteins with an N-terminal His tag and transformed into *E. coli* to produce His-HvGlpK_Ec_. Following affinity chromatography (**Figure S1**), the His-HvGlpK_Ec_ protein was further purified by SEC and found in a non-acetylated state (**Figure 3C***, lane 3*). His-HvGlpK_Ec_ was then examined as a potential substrate of HvPat2-StrepII by *in vitro* assay using acetyl-CoA as the acetyl group donor (**Figure 3C**). After incubation of these purified components, the reaction products were resolved by SDS-PAGE and analyzed by immunoblotting with anti-acetyl lysine antibodies. *Hv*GlpK purified from *H. volcanii* (*Hv*GlpK_Hv_) was used as a positive control (*lane 1*). In the absence of HvPat2, His-HvGlpK_Ec_ did not react with the anti-acetyl lysine antibody (*lanes 3 and 8*) indicating that His-HvGlpK purified from *E. coli* is not lysine acetylated. Additionally, no signal was detected that would indicate lysine acetylation of the substrate when HvPat2 and His-HvGlpK_Ec_ were incubated without acetyl-CoA (*lane 6*) meaning an acetyl-group donor is required for lysine acetylation of His-HvGlpK_Ec_. By contrast, when His-HvGlpK_Ec_ was incubated with both HvPat2 and acetyl-CoA, a strong signal for lysine acetylation of this substrate was detected (*lane 9*). A weak signal was also detected for HvPat2 (*lanes 4, 6, 7 and 9*) indicating HvPat2 is lysine acetylated, consistent with our previous finding that HvPat2 K125 is acetylated in glycerol-grown cells (34). Overall, these results suggest that the His-HvGlpK_Ec_, and therefore HvGlpK, is a substrate of HvPat2. Moreover, these results suggest that HvPat2 is a catalytically active lysine acetyltransferase of protein substrates.

### Key residues of HvPat2 predicted to be involved in acetyl-CoA binding and catalysis identified by *in silico* modeling

All GNATs form a characteristic GNAT fold that encompasses the pyrophosphate-loop, or P-loop, for acetyl-CoA binding (28). Given that there is no currently available crystal structure of HvPat2, we decided to focus our initial efforts on pinpointing the catalytic residues involved in or around the Pat2 active site by docking acetyl-CoA with the predicted Pat2 model (**Figure 4**). First, Foldseek was used to identify proteins across all domains of life that are predicted to have high structural homology with the predicted AlphaFold structure of HvPat2 (ADE05038.1)(**Figure 4A**). The search yielded both solved and predicted GNAT proteins and those with the highest structural homology (PDB: 4NXY and 2I79 and GenBank: AAG08251.1, AEW6158.1, and AN032847.1) were chosen for further comparisons. Next, the primary sequence of each protein was aligned to the HvPat2 sequence based on sequence similarity using Clustal Omega (26).

**Figure 4.**
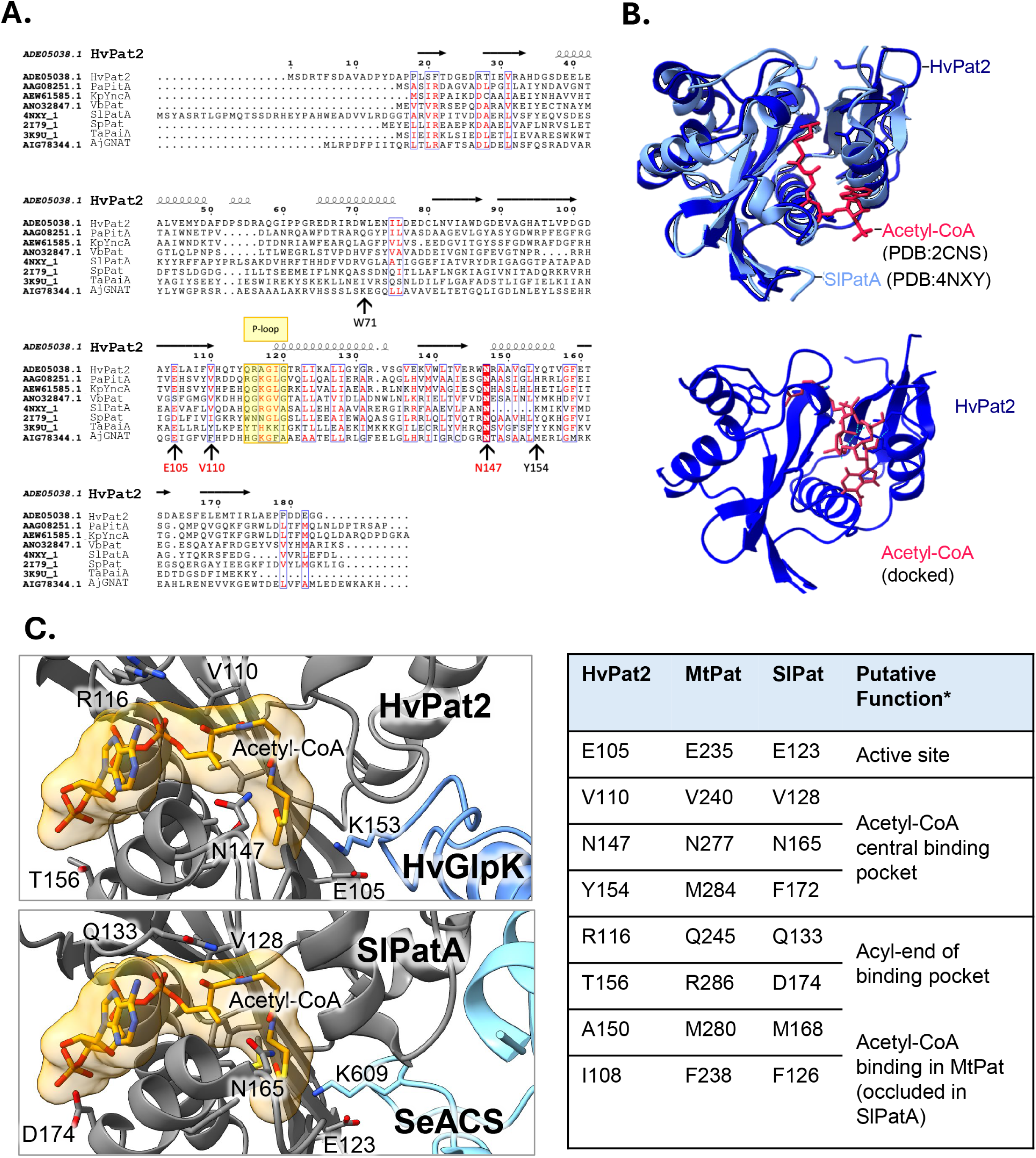
*Hv*Pat2 secondary structure prediction suggests Glu105, Val110, Asn147, and Tyr154 to be involved in A-CoA binding. **A)** Sequence conservation and secondary structure of HvPat2-related GNATs. GNAT-related lysine acetyltransferases act on a variety of substrates and share a distinct ɑ/β-fold, the conserved acetyl-coenzyme A binding domain known as the pyrophosphate binding loop (P-loop), and a catalytic Glu residue. FoldSeek was used to identify proteins with structures and sequences that are like the predicated HvPat2 structure (ADE05038). Selected sequences were aligned using ClustalW (.aln file) including (PDB or GenBank accession number): *S. lividans* SlPatA (PDB: 4NXY), *Pseudomonas aeruginosa* PaPitA (AAG08251.1), *Klebsiella pneumoniae* KpYncA (AEW61585.1), *Vibrio breoganii* VbPat (ANO32847.1), *Streptococcus pneumoniae* SpPat (PDB: 2I79), *Thermoplasma acidophilum* TaPaiA (PDB: 3K9U), *Haloferax volcanii* HvPat2 (ADE05038), and *Amycolatopsis japonica* AjGNAT (AIG78344.1). Secondary structure of the aligned sequences was predicted against SlPatA (PDB: 4NXY). Residues in red are semi-conserved and those highlighted in red are conserved. Known P-loops are boxed in yellow (24, 26). Alignment was made using ESPript 3.0 (27). **B)** 3D-structural modeling HvPat2 bound to acetyl-CoA. (*top*) ChimeraX was used to superimpose the predicted *Hv*Pat2 structure (*royal blue*) with the crystal-structure of *SI*PatA *(light blue),* and crystal structure of acetyl-CoA (from PDB: 2CNS). *(bottom)* The crystal structure of acetyl-CoA (PDB: 2CNS) was docked with *Hv*Pat2. Acetyl-CoA binding is consistent with superimposed structure and implicates Trp71, Glu105, Val110, Asn147, and Tyr154 in forming H-bonds with acetyl-CoA. This conformation represents the highest affinity docking position possible between *Hv*Pat2 and acetyl-CoA (-8.1 kcal/mol; PyRx). **C)** Docking of HvGlpK with HvPat2 bound with acetyl-CoA. AlphaFold model of HvPat2 bound to HvGlpK. While the ipTM score of 0.28 for this model is low, the site of HvGlpK acetylation (K153) (34) is in close proximity to the proposed HvPat2 active site residues. X-ray crystal structures of *Mycobacterium tuberculosis* MtPat bound to acetyl-CoA (PDB: 4avb) and *Streptomyces lividans* SlPat bound to *Salmonella enterica* SeACS (acetyl-CoA synthetase)(PDB: 4u5y) were used for comparison. Contacts of HvPat2 E150, V110, N147, and Y154 residues are highlighted. 3D-structural comparison of key residues of the bacterial Pat enzymes to HvPat2 is noted on the right.

Despite having overall low sequence homology across all GNATs, secondary structures that form the characteristic GNAT fold are highly conserved across species (10). To determine if this trend could be observed in our dataset, the multiple sequence alignment file overlaid with the HvPat2 secondary structure predicted by AlphaFold (35) using ESPript 3.0 (27) (**Figure 4A**).

The results suggest that the predicted secondary structures formed by HvPat2 match the universally conserved GNAT fold identified in previous studies (10). Additionally, the highly conserved catalytic Glu residue (E105 in HvPat2) and a plausible P-loop (Q115-G120) were observed, among other semi-conserved residues including V110 and the absolutely-conserved N147 residue (30, 36). Next, the AlphaFold-generated 3D-structural model of HvPat2 was superimposed with the X-ray crystal structure of *S. lividans* PatA (SiPatA) bound to acetyl-CoA to identify the putative acetyl-CoA binding pocket of HvPat2. To propose specific residues involved with acetyl-CoA binding, ChimeraX was used in conjunction with PyRx molecular docking software **(Figure 4B)**. The predicted 3D structure of HvPat2 was prepared for docking by removing water molecules, assigning partial charges, and adding hydrogens using ChimeraX dock prep software. Using the prepared HvPat2 and a crystal structure of acetyl-CoA (PDB: 4CNS), the most energetically efficient conformation in which acetyl-CoA binds to the enzyme within the specified binding cleft was predicted using PyRx. The PyRx molecular docking tool runs on Autodock Vina and returns the most likely conformation with the lowest binding energy. Docking results implicated five *Hv*Pat2 residues that participate in hydrogen bonding with acetyl-CoA: Trp71, Glu105, Val110, Asn147, and Tyr154. Secondary structural analysis revealed that four of the five residues are positioned close to the proposed P-loop and β-bulge regions of the enzyme (E105, V110, N147, and Y154). With this in mind, we then modeled HvPat2 docked with acetyl-CoA and HvGlpK **(Figure 4C)**. These results support prior docking models and suggest that Val110, Asn147, and Tyr154 residues appear to be involved in the central pocket for binding acetyl-CoA while E105 may function as a catalytic residue in the active site.

### Key residues identified to be important for HvPat2 function

To investigate the function of the implicated HvPat2 residues, E105, V110, N147, and Y154 were mutated to alanine using an rrPCR-based site-directed mutagenesis approach with plasmid pJAM4017 as the template, which expresses HvPat2-StrepII (25). The resulting plasmids were transformed into H26 Δ*pat2*, and strains expressing these proteins were cultured, followed by purification of the proteins using StrepII affinity chromatography as previously described for HvPat2-StrepII. Each of the HvPat2 variants purified as a single band that migrated around 25 kDa when resolved on SDS-PAGE **(Figure 5A)**. Interestingly, the HvPat2 Y154A variant was also found to co-purify with a protein migrating around 50 kDa.

**Figure 5.**
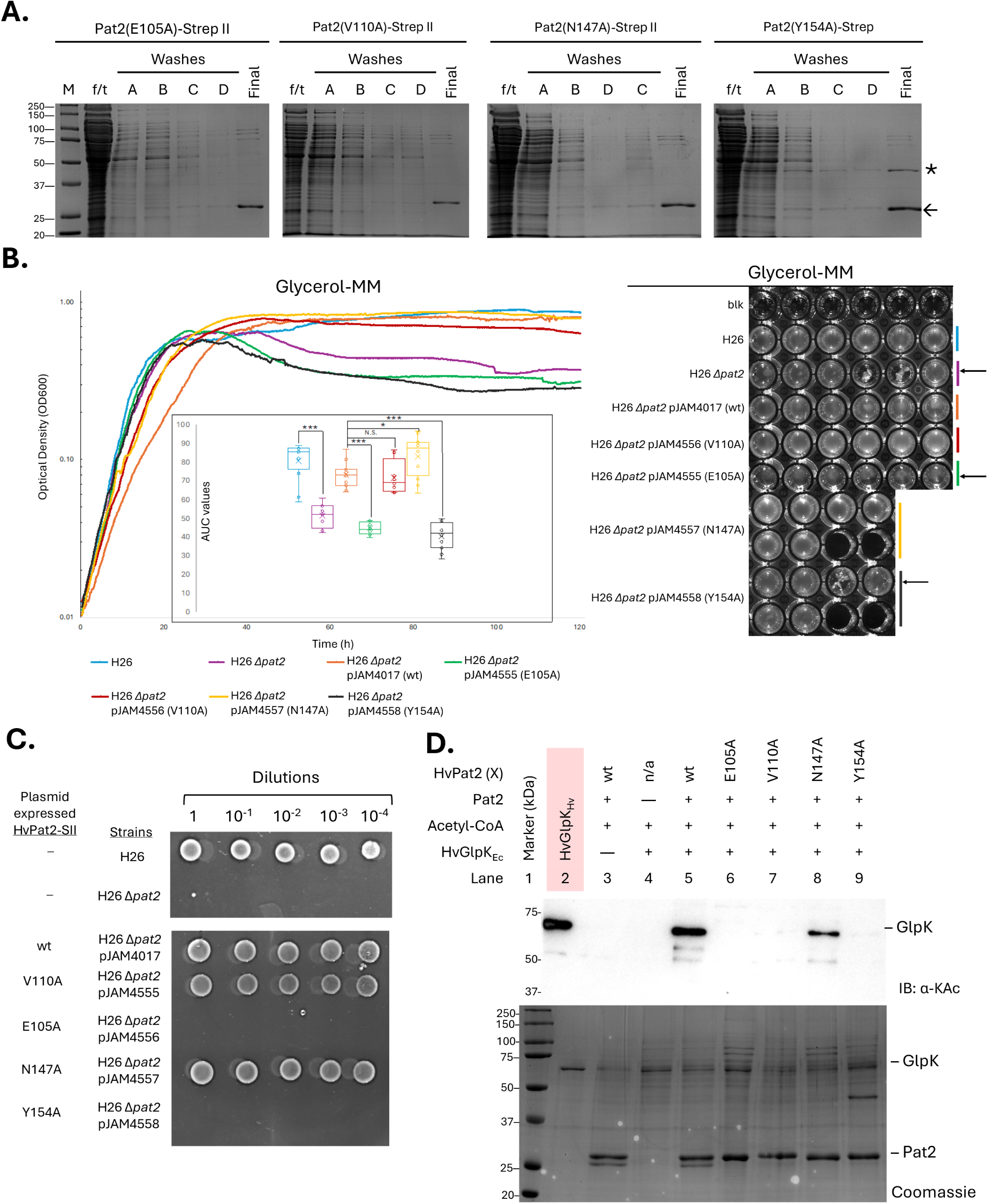
HvPat2 residues identified by *in silico* 3D-modeling found important for catalyzing lysine acetylation of HvGlpK by *in vitro* assay. A) Purification of HvPat2-StrepII variant proteins from *H. volcanii*. Strains for purification were based on H26 *Δpat2* carrying plasmids pJAM4555 (HvPat2 N147A-StrepII), pJAM4556 (HvPat2 V110A-StrepII), pJAM4557 (HvPat2 N147A-StrepII), or pJAM4558 (HvPat2 Y154A-StrepII). Cells were cultured in ATCC974. Recombinant protein expression was induced by adding 2 mM L-Trp when cultures reached an OD_600_ 0.6-0.8. Cultures were continued overnight and then harvested by centrifugation. Cell pellets (1.5 g) were lysed using a French Press. StrepII tagged proteins were purified by applying the clarified lysate to columns packed with StrepTactin resin. Flowthrough (f/t) was collected and reapplied to column for a second set of washes. StrepII tagged proteins were eluted with binding buffer supplemented with 1 mM desthiobiotin. Fractions of f/t (1 µL), first set of washes (5 µL), and final eluted protein (5 µL) were separated on 12 % SDS-PAGE. Protein gels were stained with Coomassie blue for 30 min, briefly rinsed with diH_2_O, then destained with fresh diH_2_O overnight. B) Growth curves suggest E105 and Y154 are important for HvPat2 activity during glycerol metabolism. Growth curves (*left*) of the parent H26 (light blue), H26 Δ*pat2* (purple) strains transformed with plasmids expressing Pat2 wild-type (orange) or site-directed mutants: E105A (green), V110A (red), N147A (yellow), or Y154A (black) were grown in glycerol minimal media as previously described and compared against the parent strain (H26; light blue). Inset graphs show area under the curves for each strain. * p-value ≤0.05, ** p-value <0.01, *** p-value <0.0001, N.S. no significant difference. Image of plate (*right*) immediately following growth curve demonstrates cells settled at bottom of the well and depicts the “clumping” phenotype (arrow). C) HvPat2 E105 and Y154 appear important for overcoming the loss of viability of the *Δpat2* mutant strains after growth in GMM. Cells from the GMM growth curve were serially diluted, spotted onto ATCC974 rich medium plates and incubated for one week at 42°C. The results demonstrate that the aggregated cells at bottom of wells are no longer viable. Similar results are observed on GMM plates. D) All four residues (E105, Y154, V110 and N147), predicted to be involved in catalysis or or acetyl-CoA binding, were found important for full HvPat2 activity in mediating the lysine acetylation of HvGlpK *in vitro*. Lysine acetylation reactions with HvPat2 variants were performed *in vitro* as previously described. Products were resolved by 12 % SDS-PAGE and stained with Coomassie or detected using anti-acetyl lysine antibodies.

Analysis of growth on glycerol (GMM) with the Δ*pat2* mutants expressing each of the HvPat2-StrepII variants was performed alongside the H26 parent, Δ*pat2* mutant, as well as the Δ*pat2* mutant expressing the ‘wild-type’ HvPat2-StrepII *in trans* **(Figure 5B)**. As previously observed, the Δ*pat2* mutant demonstrated a drop in OD_600_ during late stationary phase, while the parent strain and the Δ*pat2* mutant strains expressing HvPat2 V110A and N147A did not, suggesting that expression of these HvPat2 variants can complement the *pat2* deletion *in trans* for growth on glycerol. By contrast, the OD_600_ of the strains expressing HvPat2 E105A and Y154A dropped in stationary phase similar to what was observed for the Δ*pat2* mutant in GMM. Further analysis revealed a correlation between those strains that displayed a drop in OD_600_ and clumping at the bottom of the wells in the 96-well plate in a non-uniform manner. Additionally, the strains that demonstrated this clumping phenotype were no longer viable when transferred from the GMM cultures in the 96-well microtiter plate to either rich medium (ATCC974) or GMM agar plates **(Figure 5C).** These results suggest that HvPat2 E105A and Y154A are inactive, while the V110A and N147A variants can complement the *Δpat2* mutant for growth.

To further investigate residues predicted by *in silico* analysis to be important for acetyl-CoA binding or catalytic activity, the lysine acetylation activity of purified HvPat2-StrepII variants was assessed by *in vitro* assay using acetyl-CoA and HvGlpK as substrates **(Figure 5D)**. In contrast to the robust activity of wild-type HvPat2-StrepII, no catalytic activity was detected for the E105A and Y154A variants. The V110A variant showed minimal activity, while N147A exhibited reduced activity. These findings are consistent with the *in vivo* results and indicate that E105 and Y154 are critical for enzymatic function.

## Discussion

This study reveals that deletion of *pat2*, but not *pat1*, leads to impaired growth of *H. volcanii*, characterized by a carbon source-dependent clumping of cells that results in a dramatic reduction in viability during early stationary phase on glycerol. Consistent with the *Δpat2* mutation being responsible for this phenotype, this growth defect is alleviated by restoring expression of *pat2 in trans*. Through *in vitro* reconstitution, we show that HvPat2 can catalyze the lysine acetylation of glycerol kinase (HvGlpK). Given that deletion of *pat2* impairs growth on glycerol but not glucose and that HvGlpK is essential for glycerol metabolism (37), we suggest that HvPat2-mediated lysine acetylation regulates HvGlpK activity and is critical for *H. volcanii* survival on glycerol minimal medium.

To better understand HvPat2 function, *in silico* 3D structural modeling was used to guide site-directed mutagenesis, followed by analysis of the resulting HvPat2 variants. Four residues (E105, V110, N147, and Y154) were targeted for alanine-scanning mutagenesis based on their predicted role in catalysis and/ or acetyl-CoA binding. Of these, the E105A and Y154A variants were found to abolish HvPat2-mediated lysine acetylation of HvGlpK and impair cell growth on glycerol. Substitutions at V110 and N147 also reduced lysine acetylation *in vitro* but had a milder impact *in vivo*, suggesting that even low or undetectable HvPat2 activity, as measured *in vitro*, may be sufficient to maintain cell viability on glycerol. The HvPat2 Y154A variant was found to co-purify with an additional protein, suggesting it may have a higher affinity for certain proteins than the HvPat2 ‘wild-type’ and other variants. While the identity of the protein that co-purifies with HvPat2 Y154A remains to be determined, it does not appear to be lysine acetylated. One could speculate that it may function as a substrate for HvPat2 given that the Y154A mutant does not display any catalytic activity. Alternatively, if the unknown protein is not a HvPat2 substrate, it may function as a regulator of HvPat2 activity.

While most KATs in the GNAT family differ in sequence similarity and substrate specificity, they typically rely on a sequential or ping-pong catalytic mechanism (33). The ping-pong mechanism requires either a cysteine or serine to be properly oriented in the KAT active site, whereas a Glu or Asp is positioned in the active site for KATs that function through a sequential mechanism. Through this study, we can speculate that the E105 residue is positioned to acts as a general base for deprotonation of the targeted lysine residue. Through a nucleophilic attack, the lysine residue then acts on acetyl-CoA to transfer the thioester carbonyl carbon onto itself.

Lysine acetylation is suggested to play a central role in regulating carbon metabolism in *H. volcanii*, potentially through HvPat2-mediated modification of key enzymes like glycerol kinase (HvGlpK). Our prior proteomic analysis of glycerol-grown cells reveals extensive lysine acetylation across proteins involved in central carbon metabolism, including HvGlpK, one of the most highly acetylated enzymes under these conditions (**Figure 6**) (34). HvGlpK is essential for glycerol catabolism and lies at a critical metabolic junction, converting glycerol to glycerol-3-phosphate, which feeds into central metabolic pathways (38). While *H. volcanii* can metabolize multiple carbon sources, including simultaneous use of glycerol and fructose or sequential use of glycerol followed by glucose in diauxic growth (38), our findings indicate that the lysine acetyltransferase HvPat2 may be critical for regulating this metabolic flexibility. *Δpat2* mutants exhibit a severe growth defect specifically on glycerol, but not glucose, suggesting that HvPat2-mediated acetylation is essential for glycerol metabolism. This phenotype likely stems from disrupted acetylation of HvGlpK, implicating post-translational control by HvPat2 as a key determinant of carbon source utilization. Unlike bacterial systems where GlpK is regulated allosterically by fructose bisphosphate and phosphotransferase system (PTS) proteins (39, 40), *H. volcanii* appears to rely on lysine acetylation for fine-tuning GlpK activity, reflecting its distinct metabolic architecture as a heterotrophic, facultatively anaerobic archaeon capable of using a range of electron acceptors (41–43). Together, these data support a model in which HvPat2 modulates central carbon flux by acetylating HvGlpK, thereby influencing the organism’s ability to adapt to different carbon sources, particularly under glycerol-rich conditions.

**Figure 6.**
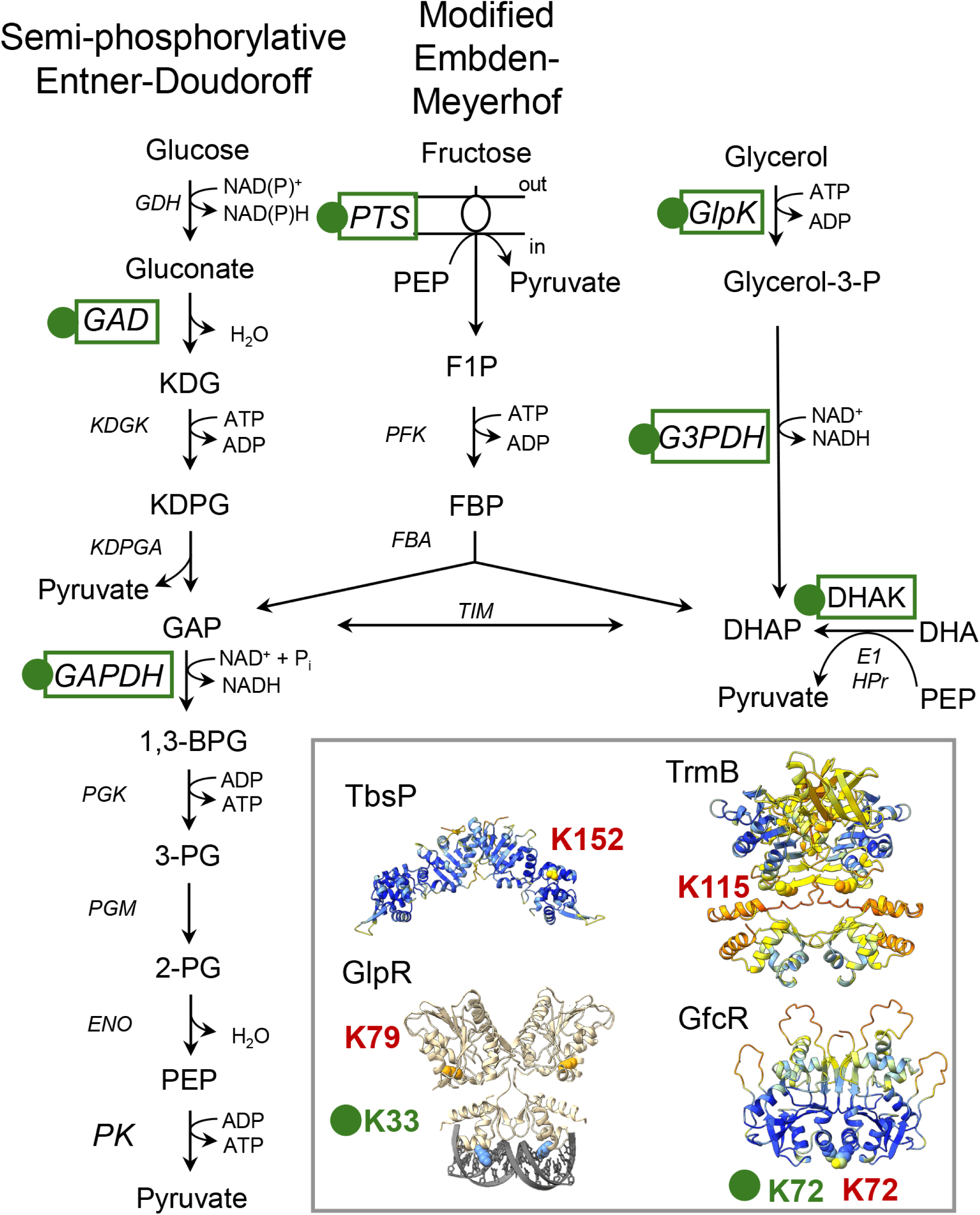
Lysine acetylation of enzymes and transcription factors associated with central carbon metabolism in *H. volcanii*. Kac, lysine acetylation. Green dot, proteins with Kac sites identified in glycerol grown *H. volcanii* based on previous study (34). Red, Kac sites of *H. mediterranei* TF homologs identified in glucose-YPC grown cells (44). AlphaFold models of transcription factors associated with central carbon metabolism with Kac sites: TbsP and TrmB (45), GfcR (46), and GlpR (37, 47, 48). Metabolites: KDG, 2-keto-3-deoxy-gluconate; KDGP, 2-keto-3-deoxygluconate 6-phosphate; GAP, glyceraldehyde 3-phosphate; 1,3-BPG, 1,3-bisphosphoglycerate; 2-PG and 3-PG, 2- and 3-phosphoglycerate; PEP, phosphoenolpyruvate; F1P, fructose-1-phosphate, FBP, fructose bisphosphate; DHAP, dihydroxyacetone phosphate; DHA, dihydroxyacetone. Enzymes: GDH, glucose dehydrogenase; GAD, gluconate dehydratase; KDGK, KDG kinase; KDGPA, KDGP aldolase; GAPDH, GAP dehydrogenase; PGK, phosphoglycerate kinase; PGM, phosphoglycerate mutase; ENO, enolase; PK, pyruvate kinase; PTS, phosphotransferase system(49), PFK, phosphofructokinase; FBA, fructose bisphosphate aldolase; TIM, triosephosphate isomerase; GlpK, glycerol kinase; G3PDH, glycerol-3-phosphate dehydrogenase; DHAK, dihydroxyacetone kinase (linked to PTS-like EI and HPr phosphorylation cascade)(50, 51). TCA cycle enzymes are all Kac modified on glycerol.

Overall, we have identified HvPat2 to function as a lysine acetyltransferase and to modify glycerol kinase (HvGlpK), an enzyme central to glycerol metabolism. Consistent with this metabolic enzyme target, we find HvPat2 to be required for survival of *H. volcanii* on glycerol, but not other carbon sources like glucose. We have shown that residues predicted to be involved in the HvPat2:acetyl-CoA:substrate binding interface play a key role in HvPat2 activity as well as the ability of cells to grow on glycerol. To expand on this work, it would be of interest to determine the crystal structure of the HvPat2 enzyme in its apo (unbound) form as well as when bound with HvGlpK to identify the complete active site of HvPat2. Understanding the molecular mechanisms used to regulate lysine acetyltransferases like HvPat2, which consist of a single GNAT domain, is also of significant interest. Whether the protein partner of HvPat2 Y154A, the UspA domain protein encoded by *hvo_1820* that overlaps the coding sequence of *pat2*, acetylation of HvPat2 K125, Sir2-mediated deacetylation, or other mechanisms are associated with this regulation remains to be determined.

## Acknowledgments

Funds awarded to JMF for this project were through the U.S. Department of Energy, Office of Basic Energy Sciences, Division of Chemical Sciences, Geosciences and Biosciences, Physical Biosciences Program (DOE DE-FG02-05ER15650) and the National Institutes of Health (NIH R01 GM57498).

